# Inflammatory Breast Cancer: a model for investigating cluster-based dissemination

**DOI:** 10.1101/119479

**Authors:** Mohit Kumar Jolly, Marcelo Boareto, Bisrat G Debeb, Nicola Aceto, Mary C Farach-Carson, Wendy A Woodward, Herbert Levine

**Affiliations:** Center for Theoretical Biological Physics and Department of Bioengineering, Rice University, Houston, TX, USA; Department for Biosystems Science and Engineering (D-BSSE), ETH Zurich, Basel, Switzerland.; Department of Radiation Oncology, The University of Texas MD Anderson Cancer Center, Houston, TX, USA; MD Anderson Morgan Welch Inflammatory Breast Cancer Research Program and Clinic, The University of Texas MD Anderson Cancer Center, Houston, TX, USA; Department of Biomedicine, Cancer Metastasis, University of Basel, Basel, Switzerland; Department of BioSciences, Rice University, Houston, TX, USA; Department of Diagnostic and Biomedical Sciences, UTHealth School of Dentistry, Houston, TX, USA

## Abstract

Metastases claim more than 90% of cancer-related patient deaths and are usually seeded by a subset of circulating tumor cells (CTCs) shed off from the primary tumor. In circulation, CTCs are found both as single cells and as clusters of cells. The clusters of CTCs, although many fewer in number, possess much higher metastatic potential as compared to that of individual CTCs. In this review, we highlight recent insights into molecular mechanisms that can enable the formation of these clusters - (a) hybrid epithelial/mesenchymal (E/M) phenotype of cells that couples their ability to migrate and adhere, and (b) intercellular communication that can spatially coordinate the cluster formation and provide survival signals to cancer cells. Building upon these molecular mechanisms, we also offer a possible mechanistic understanding of why clusters are endowed with a higher metastatic potential. Finally, we discuss the highly aggressive Inflammatory Breast Cancer (IBC) as an example of a carcinoma that can metastasize via clusters and corroborates the proposed molecular mechanisms.

## Introduction

Despite decades of advances in cancer biology, metastasis remains the primary reason for cancer-related deaths^1^. Cancer metastasis is a multistep cascade in which cancer cells escape the primary organ, enter and typically travel through the lymph and/or blood vasculature, and then exit at distant organs, eventually colonizing and proliferating at these sites leading to largely incurable stage IV disease. The metastatic cascade is highly challenging for those breakaway cells, with extremely high rates of attrition – only an estimated 0.2% of disseminated tumor cells being able to successfully seed secondary tumors or metastases^2^. Thus, the ability to initiate metastases is a key bottleneck during cancer progression and presents an ideal window for therapeutic targeting^3^.

The most well-studied mechanism proposed to facilitate metastasis is single-cell dissemination enabled by an Epithelial-to-Mesenchymal Transition (EMT). EMT is a process through which epithelial cells lose their traits of apico-basal polarity and cell-cell adhesion and gain migratory and invasive traits typical of mesenchymal cells that enable the blood-borne dissemination of carcinoma cells^4^. Conversely, after reaching a distant organ, these cells have been proposed to undergo an MET (Mesenchymal to Epithelial Transition) – a reverse of EMT – to regain their traits of cell-cell adhesion and polarity to establish metastases^5^. However, an indispensable role of EMT and MET has been called into question recently^6–8^.

Besides single-cell dissemination enabled by EMT, an alternative mechanism for metastasis that has emerged from recent studies is collective migration by clusters of Circulating Tumor Cells (CTCs). Although rare as compared to individually migrating CTCs, clusters of CTCs can individually form up to 50-times more metastases^9^. *In vivo* experiments and clinical data across multiple cancer types demonstrate that these clusters typically contain fewer than 10 cells^10,11^, clearly suggesting that clustered migration provides emergent, i.e. or ‘whole is greater than sum of its parts’, advantage for metastasis. The prognostic value of CTC clusters can be gauged by clinical observations, where patients with CTC clusters circulating in their bloodstream have significantly worse overall and progression-free survival than those in whom only individually migrating single CTCs are found^9^.

Therefore, identifying the molecular mechanisms that can form and maintain these clusters is of paramount importance in tackling metastasis. In this review, we highlight recent work that offers novel insights into mechanisms that can contribute to cluster formation and ascribe heightened metastatic potential to them. We then focus on a highly aggressive disease – Inflammatory Breast Cancer (IBC) – that forms clustered lymphatic emboli as a major means of metastasis and note several lines of evidence suggesting distant metastases also occur via clusters. IBC thus can serve as a model system to emphasize the critical role of the described molecular mechanisms in forming and stabilizing circulating CTC clusters – the primary villains of metastasis.

## Clusters of CTCs: their formation and entry into the circulation

The ability of tumor cell clusters to traverse the lung^12^ and their higher efficiency at forming metastases when injected intravenously in mice has been known for over four decades^13^. New insights into how these clusters are formed have emerged from recent lineage tracing techniques that showed that CTC clusters are not usually formed by random collisions during circulation; rather they are launched as clusters into the bloodstream from the primary tumors^9,10^ (Figure 1A). Aceto *et al*. established two differently-colored tumors on the left and right mammary fat pad of immunodeficient mice and observed that 96% of CTC clusters in the bloodstream and 92% of the lung metastatic foci were singly-colored. In contrast, when differently colored tumor cells were co-injected in the same mammary fat pad, it gave rise to multicolor clusters and metastases. Along the same lines, Cheung *et al*. injected different colored cancer cells intravenously at different times, yet rarely observed multicolored metastases, arguing for the absence of intravascular aggregation events. Consequently, CTC clusters can seed polyclonal metastases^9,10,14^, thereby contributing to intra-tumor heterogeneity at the metastatic site – a prognostic marker of poor survival across many diverse cancer types, independent of other clinical, pathologic, and molecular factors^15^.

Despite their significance, CTC clusters long have been believed to be incapable of traversing capillary-sized vessels. However, recent experiments using microfluidic devices illustrate that CTC clusters up to 20 cells can traverse constrictions with similar size to human capillary constrictions (5- to 10-μm) by rapidly reversibly reorganizing into single-file chain geometries^16^. Similar reorganization of CTC clusters obtained from patients when transplanted in zebrafish further suggest that multicellular CTC clusters can travel as a unit from the primary tumor through the circulation to distant organs to seed metastases^16^.

These striking observations are reminiscent of a study on 3-D reconstruction of serial tissue sections. Bronsert *et al.^17^* reconstructed the stromal border of various tumor types including invasive breast cancer and found little evidence of single cell migration. They concluded that cancer cell migration relies mostly, if not completely, on collective cell invasion, where invading cells retain at least partially physical cell-cell contacts. These cells that bud off from the primary tumor displayed some traits of EMT such as a morphological shift towards a spindle-like phenotype, decreased levels and membrane localization of E-cadherin, and increased nuclear levels of ZEB1, but their cell-cell adhesions were not completely lost. Therefore, these tumor buds were proposed to exhibit a ‘partial EMT’ (Epithelial-Mesenchymal Transition) or a hybrid epithelial/mesenchymal phenotype instead of a completely mesenchymal phenotype^17,18^.

**Figure 1.**
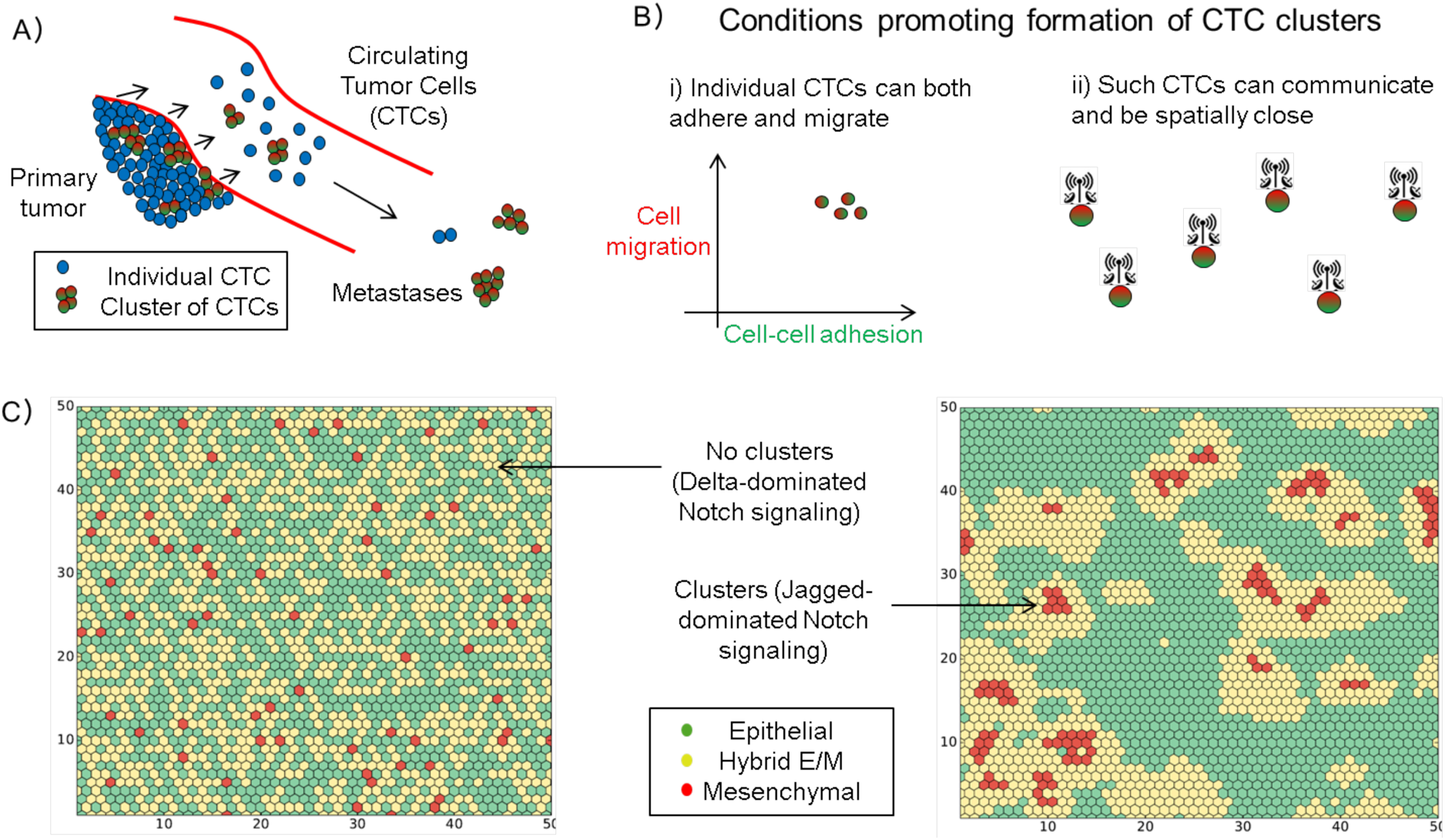
Clusters of Circulating Tumor Cells (CTCs). A) Schematic illustrating that both individual CTCs and CTC clusters can be launched from the primary tumor into the bloodstream, and clusters can form much more metastases. B) Conditions that can foster formation of CTC clusters – i) state of individual cells to be able to both migrate and adhere (i.e. hybrid epithelial/mesenchymal or E/M phenotype), and ii) cell-cell communication among these cells to spatially rearrange to form a cluster. C) Simulation of a layer of 50*50 cells that interact with one another via Notch signaling (left) predominantly via Delta ligands, and (right) predominantly via Jagged ligands. The color indicates the EMT status of each cell, as mentioned in the box (Adapted from Boareto et al. J R Soc Interface 2016 Fig 4 c,d^19^).

## Clusters of CTCs and a hybrid epithelial/mesenchymal phenotype: molecular similarities

In the context of metastasis, EMT has been defined as single-cell migration and/or invasion, along with loss of cell-cell adhesion^20^. Overexpression of transcription factors inducing EMT such as TWIST has been shown to facilitate metastasis^21^, but the role of EMT in metastasis remains controversial because a genetic knockdown of two transcription factors inducing EMT – TWIST and SNAIL-in genetically engineered mouse models (GEMM) was recently shown to be dispensable for metastasis^6^. Understanding of the role of EMT in metastasis has been confounded by two interrelated issues – (a) tacit assumption that EMT is an ‘all-or-none’ process^22^, and (b) lack of appreciation for the concept that the set of changes in cell behavior and/or lineage that have been labelled as EMT can be context-dependent. For instance, EMT during embryonic development, but not during tumor progression, refers to a lineage switch from an epithelium to a mesenchyme^23,24^. Therefore, the concept of a partial EMT or a hybrid E/M phenotype has been invoked recently to highlight that cellular plasticity during collective invasion and metastasis can be extremely fine-tuned and therefore any attempts to bin CTC clusters in a binary manner of epithelial and mesenchymal traits can be counterproductive^24–26^.

Cells in a hybrid epithelial/mesenchymal (E/M) phenotype retain at least some levels of E-cadherin – the loss of which is considered a hallmark of EMT – and co-express epithelial and mesenchymal markers and display an amalgamation of adhesion and migration to migrate collectively^24^ (Figure 1B). Such co-expressing cells have been observed in primary tumors, metastatic tumors, cell lines, mouse models and CTCs belonging to multiple cancer types^24,27–29^.

For example, H1975 cells maintain a stable hybrid E/M phenotype and migrate collectively forming finger-like projections *in vitro^29^* and adhere closely upon capture by a CTC-chip^30^. Collective migration of H1975 cells was disrupted on knockdown of GRHL2 or its downstream target OVOL2^29^. Similar roles for GRHL2 and OVOL2 have been indicated in developmental EMT^31,32^; their inhibition abrogates collective cell migration during lung morphogenesis and mammary development respectively. Thus, GRHL2 and OVOL2 can be considered as potential targets to break the CTC clusters.

Further analysis of the molecular signatures of CTC clusters and hybrid E/M phenotypes enable drawing a closer parallel between them. Cheung *et al*. demonstrated that JAG1 was one of the top differentially expressed genes in cells leading collective invasion^10^. On the other hand, Boareto *et al*. suggested that high JAG can contribute to formation of clusters of CTCs by mediating intercellular communication between cells in a hybrid E/M phenotype^19^ (Figure 1B, C). We also compared the levels of JAG and canonical epithelial (E) and mesenchymal (M) markers in individual CTCs and CTC clusters of breast cancer patients (Aceto *et al*.^9^; GSE 51827). We observed that while both individual cells and CTC clusters tend to express both E and M markers, the expression of JAG was restricted to clusters (Figure 2A). Further, co-expression of E and M markers is enriched in clusters as compared to single CTCs, and the cells expressing higher levels of JAG expressed both E and M markers (Figure 2B), thereby bolstering that targeting JAG1 can interfere with cell-cell communication among cancer and/or stromal cells^33^ and hence break CTC clusters.

**Figure 2.**
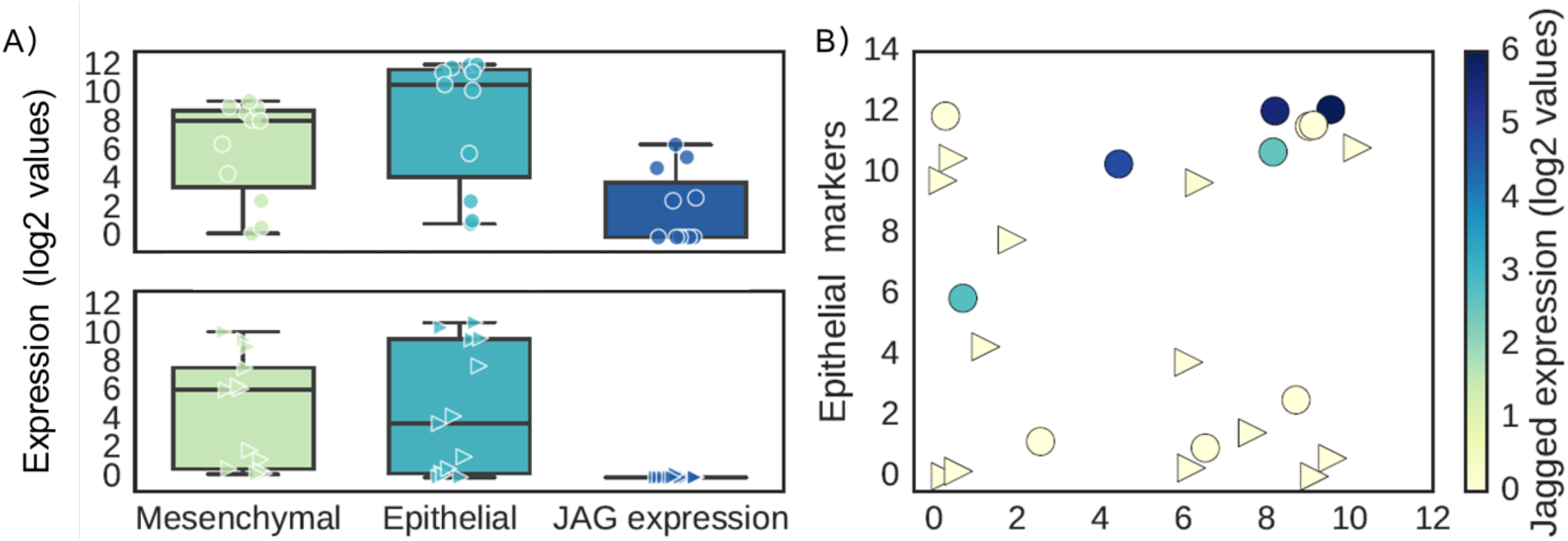
Expression pattern of 13 single CTCs and 11 CTC-clusters at single-cell level. A) Expression level of mesenchymal (ZEB1, ZEB2, SNAI1, SNAI2, TWIST1, VIM, CD44) and epithelial markers (CDH1, EPCAM, CD24) and Jagged (JAG1, JAG2) in individual CTCs (bottom) and CTC clusters (top). B) Single cells are represented based on their levels of mesenchymal markers (x-axis) and epithelial markers (y-axis). Color code represents the expression values of the ligand Jagged. Gene counts table was directly downloaded from GSE repository (Aceto et al.^9^; GSE 51827), and cells with less than 0.1 million reads were excluded from the analysis. Data was analysed via Python and Jupyter Notebook web application (source code freely available at https://github.com/mboareto/CTC_RNAseq)

Other molecular markers expressed by cancer cells and leading to collective invasion in both mouse models and human breast tumors also have been reported to induce or maintain a hybrid E/M phenotype. For instance, these invasive cells express basal differentiation markers such as P-cadherin (CDH3) and p63^34^, and knockdown of p63 is sufficient to block collective invasion. P-cadherin is a proposed marker of hybrid E/M phenotype^35^. Overexpression of the transcription factor ΔNp63α in breast cancer cells can drive collective cell migration and invasion and can induce a hybrid E/M phenotype in basal-like breast cancer (BLBC) cells by both activating miR-205 that inhibits ZEB1/2 and elevating the levels of SLUG, an activator of ZEB^36–38^ (Figure 3A, B). This coupling patterns is indicative of ΔNp63α acting as a ‘phenotypic stability factor’ (PSF) that can prevent the cells ‘that have gained partial plasticity’^31^ from undergoing a full EMT^29^ (i.e. single-cell migration and/or invasion) Additionally, ΔNp63α can trigger the secretion of matrix metalloproteinases (MMPs) to drive the invasive program^39^. P-cadherin is a downstream target of ΔNp63α^35^, and p63 gene can be activated by GRHL2^40^, another PSF^29^.

Furthermore, the expression of P-cadherin (CDH3) correlates with two PSFs - GRHL2 and its target OVOL2, and the overexpression of one or more of them can predict poor overall survival, and progression-free survival across multiple cancer types^29^. These observations are consistent with the prognostic power of a combined set of epithelial markers (cytokeratins 8 and 18) and mesenchymal markers (vimentin, fibronectin). Such co-expression, instead of the expression of mesenchymal markers solely, in invasive cancers like BLBC correlates with enhanced metastatic potential and poor survival^24^, high histologic grade, lymphovascular invasion, and can be an independent prognostic factor^41^. Cells co-expressing epithelial and mesenchymal markers are most enriched in highly aggressive cancers such as triple-negative breast cancer (TNBC) ^24,42^. Similarly, P-cadherin is aberrantly overexpressed in local advanced inflammatory breast cancer (IBC) (see below) and other highly metastatic breast cancer cells such as 4T1^43^. Put together, these results strongly suggest that a hybrid E/M phenotype can be the hallmark of collective invasion of tumors and lead to tumor aggressiveness.

It should be noted that a hybrid E/M phenotype can also be observed at a population level, i.e. in biphasic carcinomas such as carcinosarcomas that are comprised of distinct carcinomatous and sarcomatous elements components that are clonally connected^44^. An epithelial morphology of the emboli and metastases of carcinosarcomas further reinforce the idea that a partial retention of epithelial traits is critical for cells to exhibit metastatic potential^24,45–49^. However, as discussed earlier, some cancers may metastasize largely via an overt single-cell EMT-MET route, for instance, metaplastic carcinomas of pure sarcomatoid subtype may also display dismal prognosis^50^.

**Figure 3.**
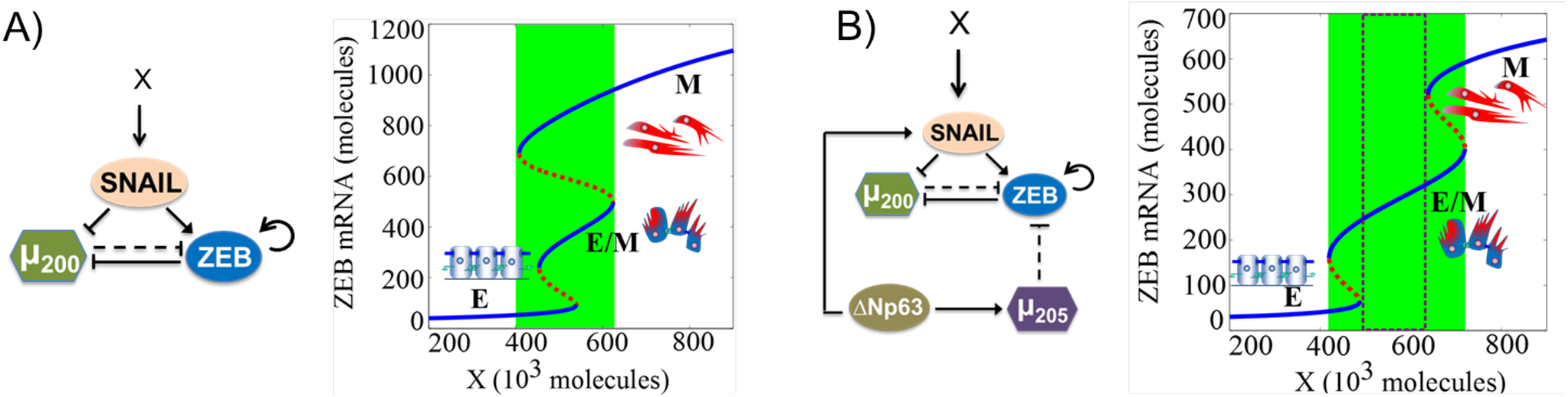
Dynamical system characteristics for miR-200/ZEB/SNAIL and miR-200/ZEB/SNAIL/ΔNp63α/miR-205 circuits. Bifurcation levels of ZEB1 mRNA in response to an external signal X driving **A)** miR-200/ZEB/SNAIL, and **B)** miR-200/ZEB/SNAIL/ΔNp63α/miR-205, representing the response of different circuits to varying levels of an EMT-inducing signal (shown as X). Solid blue curves denote stable states (phenotypes), while red dotted curves denote unstable states. Lower ZEB mRNA levels (<150 molecules) represent an epithelial (E) state, intermediate ZEB mRNA levels (~200-400 molecules) correspond to a hybrid E/M state, and higher ZEB mRNA levels (>500 molecules) denote a mesenchymal (M) state. Cartoons have been added alongside for the corresponding phenotypes. For low levels of X, cells can attain only an E state. With increasing levels of X, cells can undergo partial EMT to attain a hybrid E/M state. Further increase in X drives a complete EMT to a M state. The region marked in green represents the range of levels of X for which the hybrid E/M phenotype can exist as one of the multiple possible phenotypes; and that marked by dotted rectangle denotes the levels of X for which the hybrid E/M phenotype can exist alone. Note that the introduction of a ‘phenotypic stability factor’ (PSF) such as ΔNp63α has dramatically broadened the allowable range for a hybrid E/M phenotype. More importantly, it enabled the existence of a region where most cells can maintain a hybrid E/M state stably, i.e. a flow cytometry analysis of the region shown in dotted rectangle will identify that most cells co-express epithelial and mesenchymal markers, thereby displaying a hybrid E/M phenotype. Details of the mathematical models for both these circuits used to obtain these bifurcation diagrams are given in the Supplementary Information.

## Molecular mechanisms underlying higher metastatic potential of CTC clusters

Multiple factors can contribute to higher metastatic potential of CTC clusters, many of which tend to correlate with EMT, such as the ability to respond more effectively to mechanical signals and chemical gradients in primary tumor microenvironment as compared to individually migrating cells^51–53^, protection against apoptosis in the bloodstream upon detachment from the ECM and/or other cells^10,54^, evasion from immune attacks due to the presence of immune cells in CTC clusters and/or altered surface markers of cancer cells^11,55^, potential cooperation among the heterogeneous cell types in clusters^42^ during or before entering into circulation. Moreover, CTCs, including those in clusters, can produce numerous enzymes such as matrix metalloproteinases (MMPs)^56^ that destroy basement membrane components^39,57^, providing access to the distant tissue once lodged at different sites, therefore potentially obviating the need to extravasate actively. However, all these advantages pertain to the steps of metastatic cascade before colonization – the limiting step in metastasis formation. How do CTC clusters overcome this last and most critical bottleneck?

A key property that can explain the colonizing potential of clusters is the high tumor-initiation potential associated with a hybrid E/M phenotype, as highlighted by multiple recent studies. Grosse-Wilde *et al.^48^* segregated E, M and hybrid E/M subpopulations of HMLER cells *in vitro* and observed that hybrid E/M cells can form up to 10-times more mammospheres than either E or M cells. Ruscetti *et al.^58^* isolated hybrid E/M cells *in vivo* and demonstrated that their tumor-initiation potential was comparable to or even higher than that of the mesenchymal cells. They further illustrated that upon culturing E, hybrid E/M and M cells separately, while a majority of E and M cells retain their phenotype, more than 70% of hybrid E/M cells transition into either E or M in 24 hours, suggesting their high plasticity. Similar observations have been made *in silico^45^* and in primary ovarian cultures and tumors^59^. These results strongly bolster the emerging notion that the ‘stemness window’ or ‘tumor-initiating window’ is mostly positioned midway on the ‘EMT axis’ with E and M phenotypes as its two ends ^46,47^.

On a molecular level, hybrid E/M cells have been shown to co-express CD24 and CD44 (CD24^hi^ CD44^hi^ signature)^48^. CD24^hi^CD44^hi^ cells are present in multiple breast cancer cell lines, and their population is enriched significantly on exposure to acute chemotherapy assault, suggesting that these cells represent a drug-tolerant subpopulation capable of repopulating an entire tumor^60^. These cells have upregulated JAG1 levels but lower DLL4 levels, implicating JAG1 – already described as a key target to possible break CTC clusters – in mediating chemoresistance^19^. Furthermore, we observed that HDAC inhibitors that can induce de-differentiation of cancer cells into tumor-initiating cells and increase the mammosphere formation efficiency and ALDH activity in metaplastic SUM159 cells^61^ also expands the CD24^hi^ CD44^hi^ subpopulation (Figure 3). Put together, these studies exemplify an overlap of molecular mechanisms contributing to a hybrid E/M phenotype or CTC clusters and those mediating chemoresistance and/or metastasis-initiation.

## Inflammatory Breast Cancer (IBC): a model for clustered dissemination

Inflammatory Breast Cancer (IBC) is a highly aggressive locally advanced breast cancer with poor prognosis. In the USA, although it constitutes only 2-4% of breast cancer cases, IBC patients account for 10% of breast cancer related mortality annually. IBC patients typically have swelling and redness in skin and skin edema, instead of a mass detected by mammography^62^. At the time of diagnosis, most IBC patients already show signs of lymph node metastasis, and 30% have distant metastases, as compared to 5% of patients in non-IBC breast cancers^63^. Many molecular and behavioral aspects of IBC suggest it to be an ideal model system that manifests the traits of a hybrid E/M phenotype, collective cell invasion, and consequent aggressiveness.

**Figure 4.**
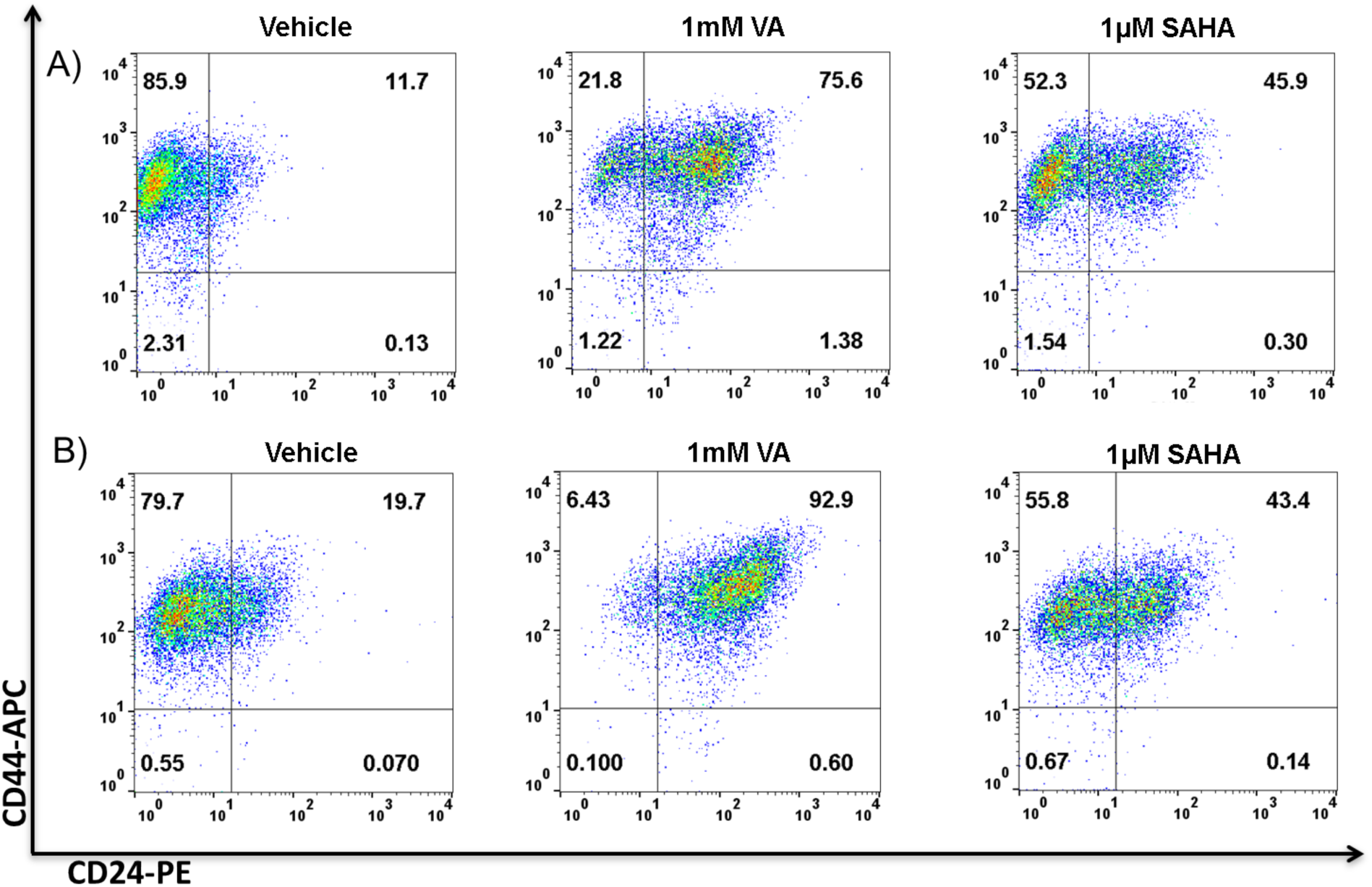
HDAC inhibitors expand the CD44^+^/CD24^+^ subpopulation. Flow cytometry analysis of expression of CD24 and CD44 in (A) ALDH^-^ and (B) ALDH^+^ SUM159 cells showing a shift from the CD44^+^/CD24^-^ in the vehicle-treated to a CD44^+^/CD24^+^ population in 1mM VA or 1 μM SAHA treated cells. Representative flow cytometry data is shown. Gating was set to unstained control cells^61^. For analyzing the CD44/CD24 subpopulation, cells were harvested with trypsin, centrifuged and re-suspended in phosphate buffered saline (PBS) (10^5^ cells/ml). Cells were incubated with APC-conjugated CD44 and PE-conjugated CD24 antibodies (BD Biosciences, San Diego, CA, USA) for 30 minutes in ice at concentrations recommended by the manufacturer. Cells incubated in PBS and cells single-stained with APC- or PE-conjugated antibodies served as controls. Cell analysis for the expression of CD44 and CD24 was performed using a Beckman Coulter machine and the data files were analyzed using FlowJo software (Treestar, Ashland, OR).

A key difference in IBC and non-IBC is the ubiquitous presence of E-cadherin in primary tumors, tumor emboli or clusters in the lymphatic system, and metastases^64,65^, a presence which might appear paradoxical given the established role of E-cadherin as a metastasis suppressor in a variety of cancers^66^. On the contrary, E-cadherin appears to augment the invasion of SUM 149 cells, an IBC cell line, by increasing the levels of matrix metalloprotease enzymes such as MMP-1 and MMP-9^66^, suggesting that E-cadherin can promote IBC progression. Recent characterization of metastatic breast cancer cells show that they retain some levels of epithelial adhesion genes such as E-cadherin^10,67^ and that E-cadherin is essential for leveraging the advantages of the osteogenic niche and consequent colonization of bone by breast cancer cells, thereby providing potential mechanistic insights into the role of E-cadherin in IBC ^68^.

Another hallmark of IBC is the presence of numerous cohesive clusters in the lymphatics and their resistance to multiple therapies^66^. As compared to non-IBC patients, IBC patients can have larger and a higher frequency of clusters of CTCs and these clusters have a stronger association with survival^69^, potentially due to their resistance to severe therapeutic assaults^70^. These clusters or emboli have accumulation of E-cadherin due to its altered trafficking^71^, accumulation which may aid the passive dissemination of these emboli^72^. Passive dissemination can facilitate metastasis in various contexts^73^, and investigating active vs. passive dissemination mechanisms during intravasation can help reconcile the controversy on the role of EMT in metastasis^74^.

Transfection of dominant negative E-cadherin in MARY-X, a mouse model for IBC that exhibits tight aggregates of individual tumor cells held by E-cadherin in suspension, reduces the formation of these emboli^75^, indicating a role for E-cadherin in maintaining the clustered phenotype. These results are reminiscent of implications of E-cadherin in collective chemotaxis *in vivo^76^*, and that knockdown of GRHL2 or OVOL2 – top activators of E-cadherin^77^ – disrupt collective finger-like motion *in vitro* in lung cancer cells^29^. Another potential mechanism through which E-cadherin can drive aggressive behavior is the survival of clusters in the bloodstream by ‘synoikis’, i.e. activation of survival signals through junctional adhesions between neighboring cells^78^.

Besides maintaining E-cadherin levels, IBC cells often also express mesenchymal proteins such as vimentin, thereby adding to the interpretation of IBC as a manifestation of a hybrid E/M phenotype^79^ – (a) FC-IBC-02 cells express vimentin alongside E-cadherin and other markers of epithelial phenotype such as EpCAM (Epithelial cell adhesion molecule), (b) compared to MDA-MB-231, multiple IBC cell lines – MDA-IBC-3, SUM 190, FC-IBC-02 and SUM149 – have intermediate levels of ZEB1, a proposed marker for hybrid E/M phenotype^80^ (c) FC-IBC-02 cells both in mammospheres and adherent conditions express SLUG (SNAI2), a key mediator of a partial EMT state during mammary morphogenesis^81^, and JAG1 that can contribute in maintaining a cluster of cells in a hybrid E/M phenotype^19^. Furthermore, FACS analysis of cell line and mouse models of IBC – SUM 149, Mary-X, FC-IBC-01, and FC-IBC-02 – indicates that a large percentage of cells are CD24^hi^ CD44^hi82^, the proposed signature for a hybrid E/M phenotype^24,48^. In contrast, mesenchymal cells such as MDA-MB-231 predominantly express CD44 (a mesenchymal stem cell marker) but lack CD24 (an epithelial marker)^82^.

The predominance of CD24^hi^ CD44^hi^ cells can be an important underlying reason for resistance of IBC against chemotherapy and radiotherapy. CD24^hi^ CD44^hi^ cells represent the adaptive drug tolerant population that is enriched upon treatment of multiple breast cancer cell lines with docetaxel^60^. These cells indicate a higher proclivity for Notch-Jagged signaling instead of Notch-Delta signaling^19^, and are consistent with observations that drug-resistant small cell lung cancer H69-AR cells have higher levels of JAG1 but lower levels of DLL4 as compared to the parental H69 population^83^. Significantly high levels of IL-6 in serum of IBC patients as compared to non-IBC patients, overexpression of IL-6 in IBC carcinoma tissues, and secretion of IL-6 by IBC cell lines SUM149 and SUM190^84,85^ and supporting stromal cells^86^ can augment Notch-Jagged signaling, thereby contributing to radiation resistance of these cell lines ^87^ and IBC progression. Notch-Jagged signaling can also mediate high tumor-initiating potential in IBC. Lymphovascular emboli of MARY-X shows an addiction for Notch 3, and its knockdown can induce apoptosis and inhibit the levels of a stem cell marker CD133. The emboli of human IBC exhibits immunoreactivity for both cleaved Notch 3 intracellular domain (ICD) and stem cell markers such as ALDH1^88^.

Multiple lines of evidence indicate that an induced transition from a hybrid E/M profile of IBC to a fully mesenchymal phenotype can reduce IBC aggressiveness. For instance, overexpression of ZEB1 or knockdown of E-cadherin, either of which can induce and often maintain a complete EMT phenotype^80,89^, reduced the *in vivo* growth of SUM 149 primary and metastatic tumors^90^. E-cadherin knockdown also reduced the *in vivo* growth of 4T1 and MARY-X cells^90^. Similarly, invasion of IBC cells is disrupted by exposure to TGFβ^91^, a potent inducer of EMT that can induce a complete EMT ^89^ and consequently single-cell migration phenotype^92^. Consistently, a common feature of multiple pre-clinical models of IBC, independent of subtype, is the expression of SMAD6, a repressor of TGFβ signaling by its ability to inhibit SMAD4^82^. Furthermore, high incidence of brain metastasis as reported in IBC patients^63^ can be reduced by knockdown of miR-141 – a miR-200 family member that prevents EMT induction suggesting that a complete mesenchymal phenotype lacks the potential to colonize the brain^93^.

## Conclusion

Overall, observations in IBC mouse and cell line models and *in vivo* experiments on CTC clusters challenge the hypothesis that a total loss of E-cadherin is necessary for metastasis. Such loss of epithelial markers has often been considered synonymous of an EMT, therefore these results question the indispensability of at least a complete abrogation of epithelial traits, and rather strongly suggest a potentially crucial role of partial retention of epithelial traits (hybrid E/M phenotype) in establishing metastasis, at least in IBC. For instance, limited E-cadherin levels at the adherens junctions of cells in a CTC cluster can orchestrate synoikis and therefore prevent CTC clusters from death in circulation.

